# Genome-wide association study of heat stress response in *Bos taurus*

**DOI:** 10.1101/2023.06.05.543663

**Authors:** Bartosz Czech, Yachun Wang

## Abstract

Heat stress is a major challenge in cattle production, affecting animal welfare, productivity, and the economic viability of the industry. In this study, we conducted a genome-wide association study to identify genetic markers associated with heat stress tolerance in Chinese Holstein cattle. We genotyped 68 cows using Illumina 150K Bovine BeadChip microarray and analyzed 112,081 single nucleotide polymorphisms using a linear model-based GWAS approach. We identified 17 SNPs distributed over three chromosomes that showed statistically significant associations with heat stress tolerance in Chinese Holstein cattle. Five of them were located in introns of two genes, *PDZRN4* and *PRKG1. PDZRN4* is involved in protein degradation pathways, while *PRKG1* encodes a protein kinase involved in smooth muscle relaxation and blood vessel dilation. Our findings highlight the potential importance of *PDZRN4* and *PRKG1* in heat stress tolerance in cattle and provide valuable genetic markers for further research and breeding programs aimed at improving heat stress tolerance in Holstein cattle. However, further studies are needed to elucidate the exact mechanisms by which these SNPs contribute to heat stress tolerance and their potential implications for practical cattle breeding strategies.

## Introduction

Heat stress in cattle occurs when the animal’s body temperature rises above physiologically normal levels due to the exposure to high temperatures, humidity, and solar radiation [1]. This can occur in both dairy and beef cattle and is a significant problem for the livestock industry, particularly in regions with hot and humid climates [2].

Heat stress can affect cattle in several ways. Firstly, it can lead to a decrease in feed intake and subsequently reduce weight gain or milk production [3]. Secondly, heat stress can result in respiratory distress, panting, and increased water consumption, which can put additional strain on the animal’s cardiovascular system [4]. Finally, severe heat stress can lead to dehydration, electrolyte imbalances, and to dramatic changes in the animal’s physiology [5]. In addition to the direct impact on the animal’s health, heat stress also results in economic losses due to reduced milk production and lower reproduction rate [6].

Exposing cattle to prolonged heat stress triggers complex molecular responses involving changes in gene activity and epigenetic modifications. This means certain genes become more or less active, and the tags on the DNA change, affecting how genes work without altering the DNA sequence itself. These changes do not just impact the current generation of cattle; they can also affect the genetic makeup of future generations. Research suggests that environmental stressors experienced by parents can leave lasting marks on their offspring’s genes. Understanding how heat stress affects genes and epigenetic markers is essential for developing strategies to help cattle cope with hot conditions and maintain their health and productivity. It also sheds light on how animals adapt to different environments over time [7].

Studies have shown that heat stress can lead to changes in the expression of genes related to immune function, metabolism, and reproduction. For instance, when cows experience heat stress, it can lead to a decrease in the activity of genes responsible for milk production, such as those encoding caseins, which are important milk proteins. Conversely, there is often an increase in the activity of genes involved in stress responses, including antiapoptotic genes like HSPA1A and BCL2. These genes likely play a role in protecting the cells from heat-induced damage. It’s interesting to note that despite these changes, the mRNA levels of genes coding for caseins, crucial components of milk, remain unaffected by heat stress. This suggests a selective response of gene expression to heat stress, prioritizing cellular protection mechanisms over milk production [8].

In the case of heat stress in cattle, GWAS can be used to identify genetic variants that are associated with e.g. body temperature, drooling score, and respiratory score. That will allow to develop breeding strategies to select animals that are more tolerant to heat stress and maintain productivity under hot and humid conditions [9].

Moreover, from the scientific perspective, GWAS allows understanding of the genetic basis of heat stress, including the biological pathways and mechanisms involved in heat stress response. Overall, the combination of NGS, genotyping microarrays, and GWAS can provide a powerful approach for the identification of genetic variants and even candidate genes associated with heat stress response in cattle. This knowledge can be used to develop new management practices, breeding strategies, and therapeutics to improve animal welfare and productivity under changing environment.

The aim of this study was to identify genetic variants associated with heat stress response in cattle that lead to a better understanding of the functional basis of heat stress tolerance in cattle.

## Materials and methods

The material consists of 68 cows representing Chinese Holstein cattle. These individuals were genotyped using the Illumina 150K Bovine BeadChip (Illumina Inc., San Diego, CA, USA), which consists of 123 268 single-nucleotide polymorphisms (SNPs). For all the animals heat stress response phenotypes were expressed by rectal temperature (RT), drooling score (DS), and respiratory score (RS). These phenotypes were represented by deregressed proofs (DRP) of estimated breeding values (EBVs) predicted as previously described in Czech, *et al*. [10] [11].

For comprehensive details regarding the measured phenotypes and sampling methodology, please refer to Czech et al. [10] [11].

Pre-processing of genotype data comprised retaining: i) individuals with call rate greater than 0.9, ii) SNPs with minor allele frequency (MAF) greater than 0.05, and iii) SNPs that were in Hardy-Weinberg equilibrium (*P* -value > 0.05). The filtration process was performed using the PLINK software (v1.90b6.21) [12]. Followingly, GWAS was performed separately for each phenotype, using the following model:

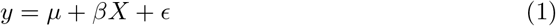

where *y* is a vector of DRPs, *X* contains SNP genotype coded as 0, 1, or 2, representing the number of reference alleles, *β* is the SNP additive effect, and *ϵ* represents residuals. The significance of a SNP effect was tested using the likelihood ratio test with the reduced model represented by model 1 without the SNP effect. The estimation of the model parameter and testing of the significance of the SNP effect were performed using the GEMMA software [13]. Additionally, the genomic inflation factor (λ) was calculated to assess the extent of inflation in the test statistics. The genomic inflation factor is a measure that compares the median of the observed test statistics to the median of the expected test statistics under the null hypothesis. It was calculated using the following formula:

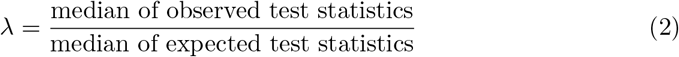

To visualize GWAS results, the qqman [14] package was used.

To control for multiple testing *P* -values were adjusted using the Bonferroni correction. Significant SNPs were considered based on the adjusted *P* -values lower than 0.05. All significant SNPs were annotated using the Variant Effect Predictor (VEP) implemented in the ensemblVEP R package with Ensembl Release 109 (Feb 2023) [15]. Additionally, the Animal QTL database was used (QTLdb) to explain the genetic basis of variation in heat stress phenotypes [16]. Next, the Gene-Set Enrichment Analysis (GSEA) was performed to detect potential functional pathways underlying the heat stress response by applying one-sided version of Fisher’s exact test. The Gene Ontology (GO) [17] and the Kyoto Encyclopedia of Genes and Genomes (KEGG) [18] were considered in GSEA implemented in the clusterProfiler R package [19].

## Results

### Genome-wide association study

The filtration process retained 112 081 out of 123 268 (91%) SNPs for GWAS for all 68 individuals. As a result for rectal temperature, 17 significant SNPs were identified, while no significant hits were observed for drooling and respiratory scores. Significant SNPs associated with rectal temperature were located on chromosomes 5, 17, and 26. On the *Bos taurus* autosome (BTA) 5 there were three significant SNPs, on the BTA17 there were 12 significant SNPs, while on the BTA26 there were only two significant SNPs. Manhattan plots were presented on Figure 1 (for rectal temperature), on the Figure 3 (for drooling score), and on the Figure 5 (for respiratory score). Quantile-quantile (QQ) plots were presented together with the genomic inflation factor on Figure 2 (for rectal temperature), on the Figure 4 (for drooling score), and on the Figure 6 (for respiratory score). Table 1 shows detailed information regarding significantly associated SNPs with rectal temperature. All SNPs were further annotated and processed through GSEA.

**Table 1.**
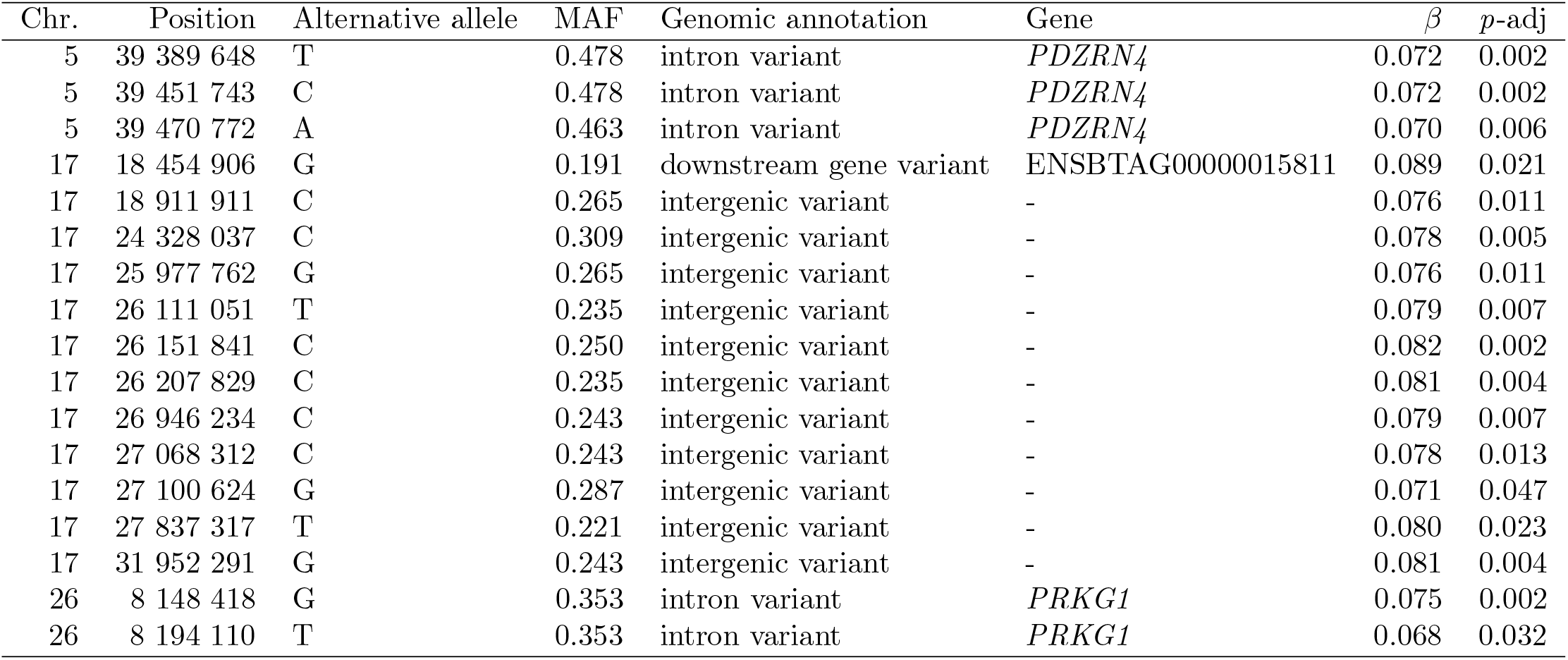
The genomic annotation of associated SNPs with rectal temperature.

**Fig 1.**
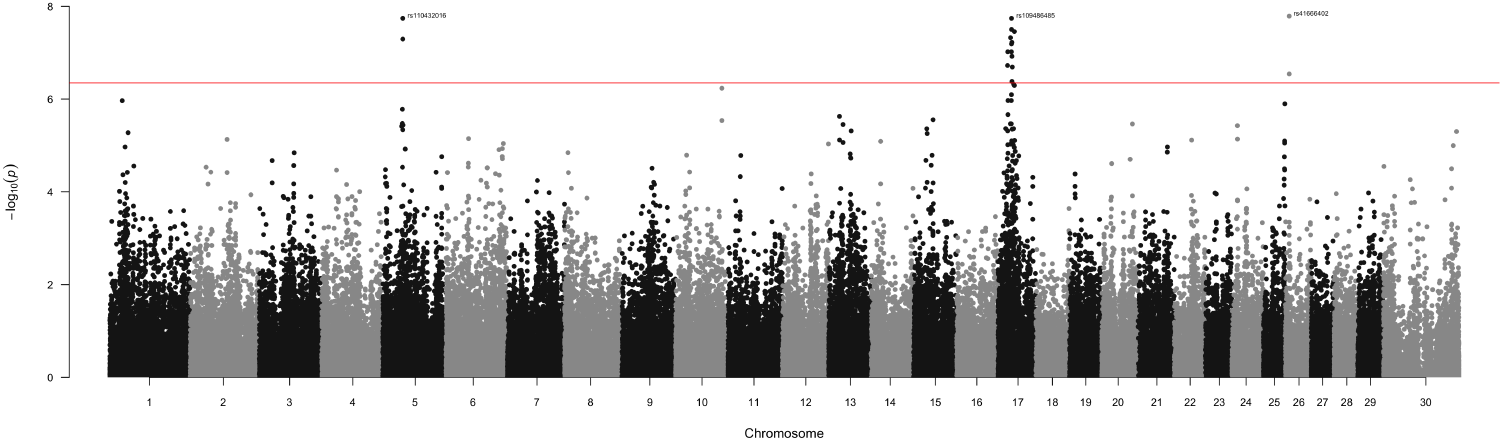
Manhattan plot for associations of SNPs with the rectal temperature. X-axis: SNPs positions on chromosomes, Y-axis: -log10 *p*-value. The red line indicates the 0.05 significance threshold corrected for multiple testing. Labels show the top significant SNPs.

**Fig 2.**
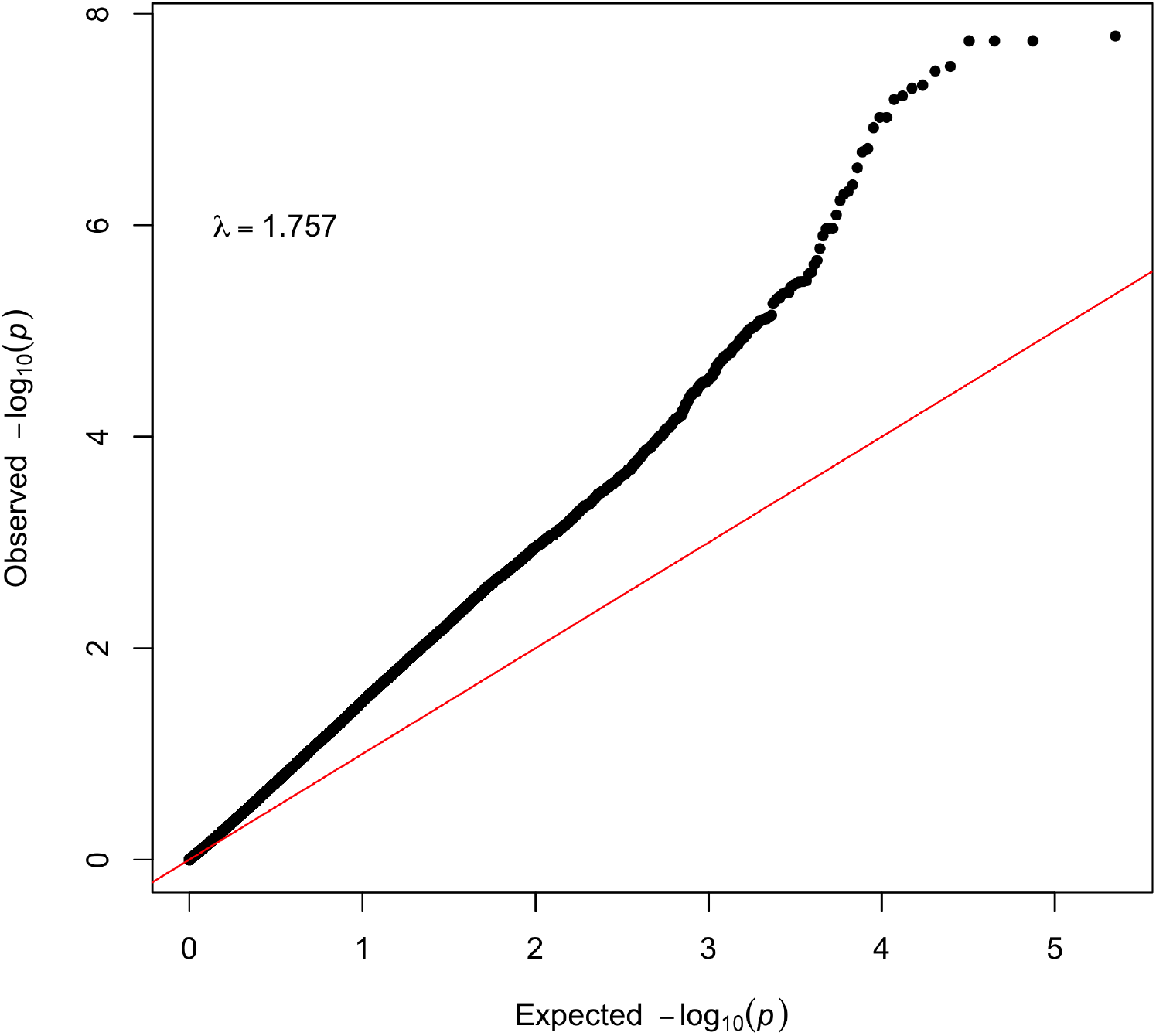
Quantile-quantile (Q-Q) plot of observed versus expected *p*-values of the GWAS results for rectal temperature. The straight line in the Q-Q plot indicates the distribution of SNPs under the null hypothesis. On the top left corner the genomic inflation factor is presented.

**Fig 3.**
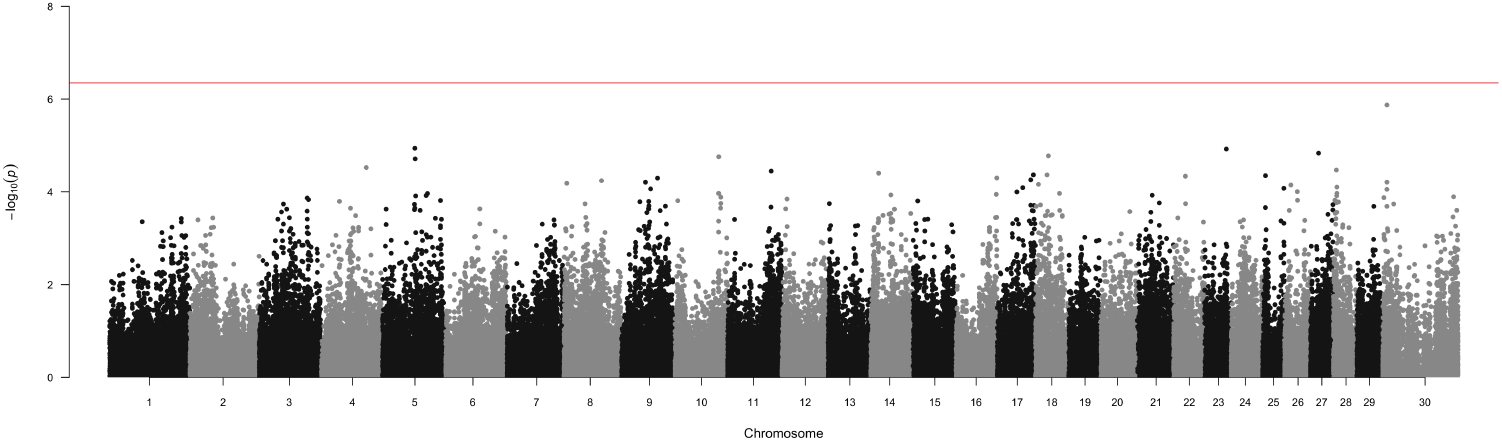
Manhattan plot for associations of SNPs with the drooling score. X-axis: SNPs positions on chromosomes, Y-axis: -log10 *p*-value. The red line indicates the 0.05 significance threshold corrected for multiple testing. Labels show the top significant SNPs.

**Fig 4.**
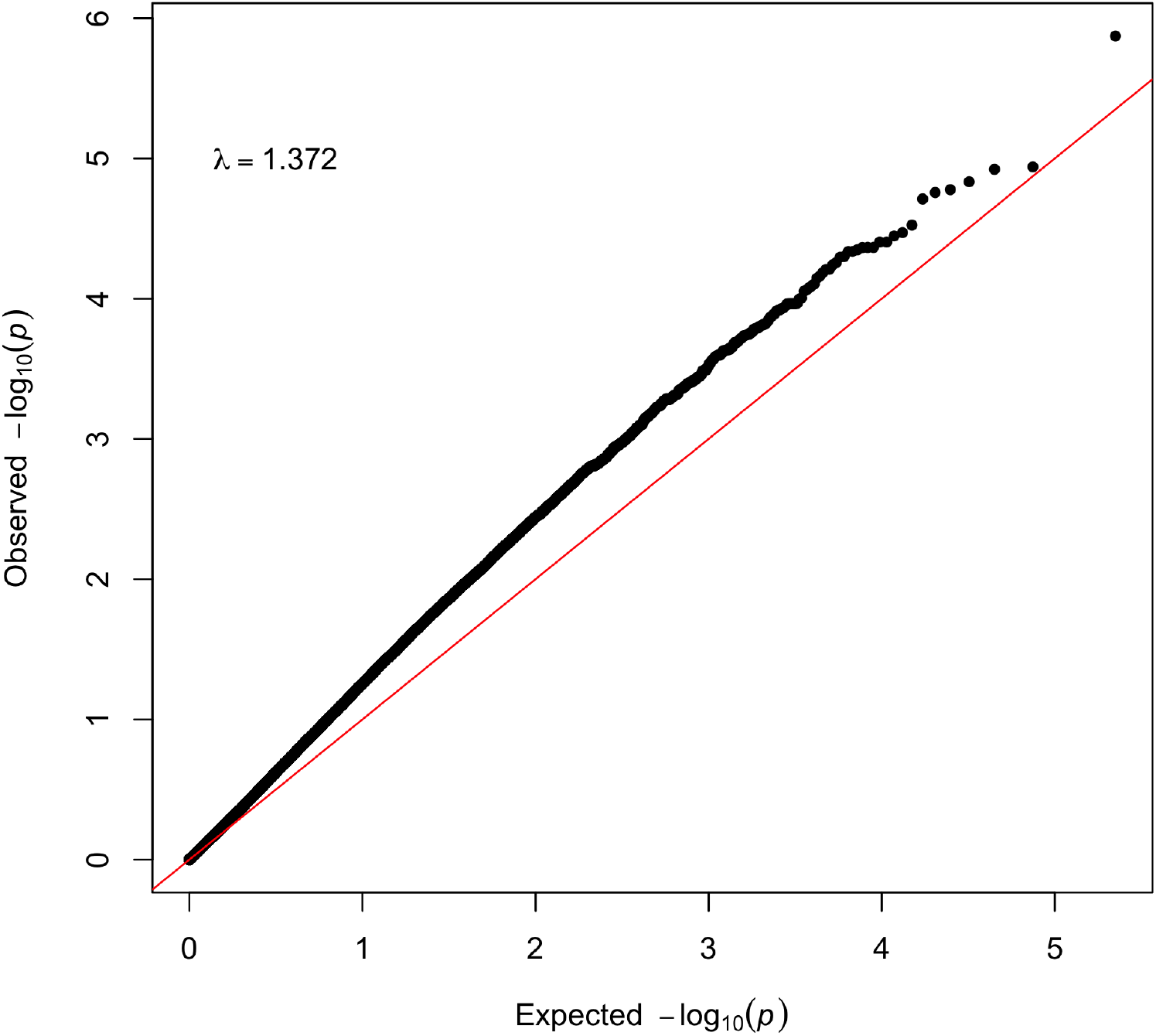
Quantile-quantile (Q-Q) plot of observed versus expected *p*-values of the GWAS results for the drooling score. The straight line in the Q-Q plot indicates the distribution of SNPs under the null hypothesis. On the top left corner the genomic inflation factor is presented.

**Fig 5.**
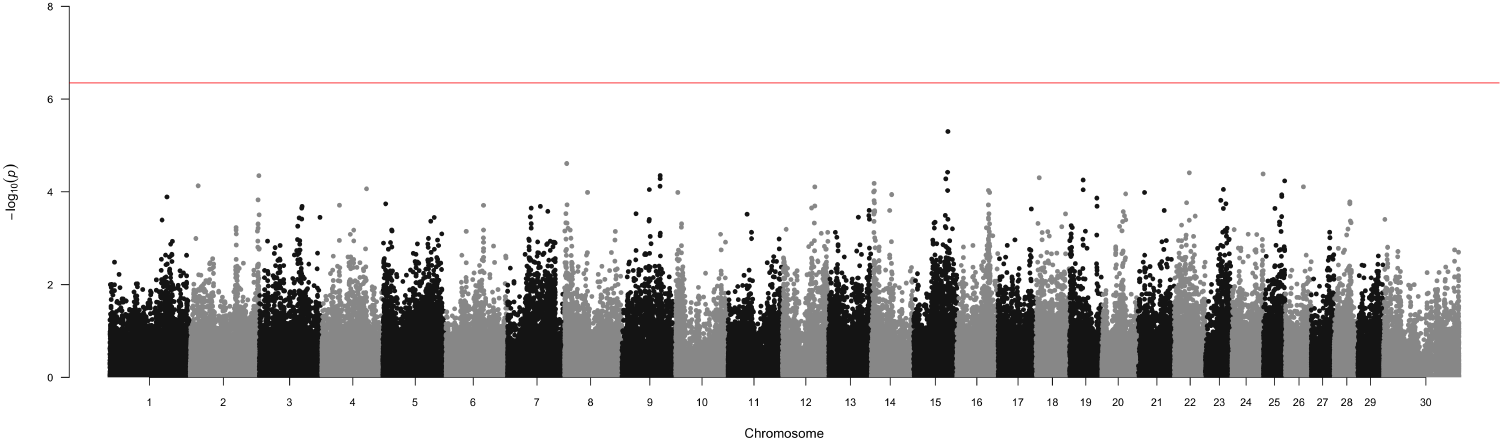
Manhattan plot for associations of SNPs with the respiratory score. X-axis: SNPs positions on chromosomes, Y-axis: -log10 *p*-value. The red line indicates the 0.05 significance threshold corrected for multiple testing. Labels show the top significant SNPs.

**Fig 6.**
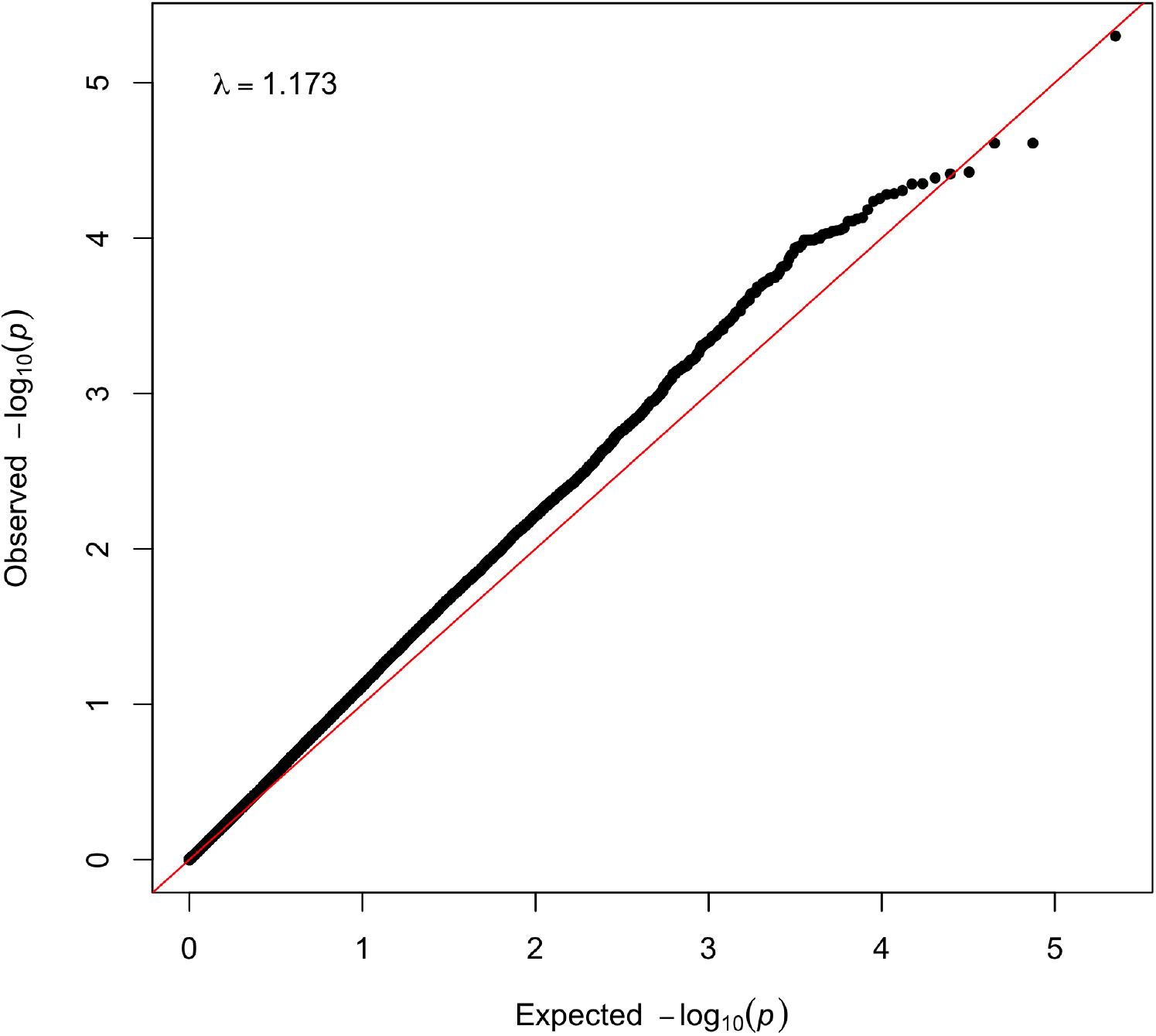
Quantile-quantile (Q-Q) plot of observed versus expected *p*-values of the GWAS results for the respiratory score. The straight line in the Q-Q plot indicates the distribution of SNPs under the null hypothesis. On the top left corner the genomic inflation factor is presented.

### Variants annotation and Gene-Set Enrichment Analysis

Results of the annotation process performed using the VEP were summarised in Table 1 which shows that all the significantly associated SNPs with rectal temperature on BTA5 and BTA26 were located in introns of *PDZRN4* (BTA5) and *PRKG1* (BTA25) genes. On BTA17, 11 out of 12 SNPs were located in the intergenic regions, while one SNP was located in the downstream part of the ENSBTAG00000015811 gene. Phenotype annotation based on the QTLdb demonstrated that SNPs 17:27837317 (rs110432016) and 17:25977762 (rs109962820) were related to a maternal component of calving ease, dairy form, daughter pregnancy rate, foot angle, milk fat percentage, milk fat yield, net merit, length of productive life, milk protein percentage, milk protein yield, rear leg placement, and teat length. GSEA based on the GO indicated that genes related to the associated SNPs were enriched in the following ontologies: GO:0003682:chromatin binding, GO:0004672:protein kinase activity, GO:0004674:protein serine/threonine kinase activity, GO:0004692:cGMP-dependent protein kinase activity, GO:0005524:ATP binding, GO:0006468:protein phosphorylation, GO:0042802:identical protein binding, GO:0005515:protein binding, and GO:0046872:metal ion binding. GSEA for the KEGG pathways showed only the enrichment of the cGMP-PKG signaling pathway (bta04022).

## Discussion

Heat stress is a major environmental challenge for livestock production, including cattle, as it can negatively impact animal health, welfare, and productivity. In this study, we performed a GWAS to identify genetic markers associated with heat stress in cattle. Our findings revealed significant associations between heat stress and single nucleotide polymorphisms (SNPs) located in the *PDZRN4* and *PRKG1* genes, shedding light on mechanisms underlying the heat stress response in cattle. This GWAS study serves as a follow-up to previous analyses of differential gene expression and differential abundance of microbiota in the context of heat stress, providing further insights into the complex interactions between genetics, gene expression, microbiota, and heat stress response. The previous studies already demonstrated the importance of rectal temperature as a main indicator of heat stress in cattle. It has been shown that of all three phenotypes (rectal temperature, drooling score, and respiratory score), rectal temperature showed a major association with gene expression and abundance of microbiota in cattle under heat stress conditions [10] [11]. Heat stress phenotype is difficult to quantify and out of the three measurements that were available in this study, only rectal temperature appeared to be the most representative of heat stress.

There are many publications on the GWASs related to heat stress in cattle, however, almost all of them focused on the standard case-control experimental design in which one cannot identify potential candidate genes responsible for the heat stress response. It is due to the complex nature of heat stress and the involvement of multiple genes and environmental factors. However, unlike previous studies that have focused on controlled experimental environments, our study examined animals in their production environment, which provides valuable insights into the genetic factors influencing heat stress tolerance in cattle under production conditions. While this approach allows for capturing the genetic variation present in the population, it also has limitations, including potential confounding factors and the lack of control over environmental variables that may interact with the heat stress phenotype in real production systems.

The identification of SNPs in *PDZRN4* and *PRKG1* associated with heat stress in cattle suggests that these genes may play a role in the physiological response of cattle to heat stress. *PDZRN4* gene (PDZ domain containing ring finger 4) also known as *LNX4* (Ligand of Numb Protein-X 4) plays a potential role as a tumor suppressor gene and may have an antiproliferative effect on hepatocellular carcinoma cell proliferation [20]. Another study showed that *PDZRN4* is a functional suppressor of prostate cancer growth [21]. Nevertheless, there are no studies in which *PDZRN4* was indicated as a candidate gene related to the heat stress response. Studies related to other livestock species showed, that this gene might affect the fat metabolism in pigs [22]. Moreover, *PDZRN4* was found as a significant gene associated with poor sperm motility in Holstein-Friesian bulls [23]. However, another gene identified in this study was *PRKG1* that encodes a protein called cGMP-dependent protein kinase 1 [24]. *PRKG1* was found as the associated gene with tick resistance in South African Nguni cattle [25]. Another study showed the importance of this gene in the local adaptation of indigenous Ugandan cattle to East Coast Fever [26]. However, the most interesting is that gene *PRKG1* was already found as a gene with a key role in body thermoregulation. In the study focused on the cold adaptation of the indigenous Siberian populations, *PRKG1* has been shown as the gene involved in cold acclimatization [27]. Another study showed that this gene is key to minimizing heat loss by regulating blood vessel constriction in Yakutian horses [28]. There is also a study confirming the important role of *PRKG1* in temperature regulation in a cold environment in Amur tiger [29]. Regarding the heat stress phenomenon, it has been shown, that *PRKG1* is associated with the heat stress adaptation in Egyptian sheep breeds [30].

Heat stress tolerance is a complex trait that involves the interplay of multiple genetic and environmental factors [31]. The SNPs located in *PDZRN4* and *PRKG1* provide valuable markers for selecting heat stress tolerant animals in breeding programs. This may lead to the development of genomic selection programmes towards improved heat stress resistance in cattle and improve animal welfare and productivity in hot climates.

### Study limitations

It is important to note that our study has some limitations. First, the sample size may affect the statistical power to detect all SNPs associated with heat stress. Small sample sizes make it challenging to demonstrate deviation from Hardy-Weinberg equilibrium, as the limited statistical power can obscure potential deviations and lead to some problematic genotypes going undetected. Despite this limitation, research based on smaller sample sizes can still provide valuable scientific insights, especially for complex phenotypes where obtaining large sample sizes is often impractical.

The QQ plots generated for analyses showed that the black line, representing hte observed *p*-values, was above the red line, which represents the expected *p*-values under the null hypothesis. The genomic inflation factors (λ) were calculated to quantify this deciation and were found to be 1.76, 1.37, and 1.18 for the three analyses conducted. Theses λ values indicate moderate inflation, suggesting that there may be some underlying factors contributing to the inflation, such as population stratification or cryptic relatedness. However, it is important to note that our study had a relatively small sample size, which can also contribute to the observed inflation. Small sample sizes are more susceptible to random variation, leading to an overestimation of the test statistics and, consequently, higher λ values. Despite the sligh elevation in the λ values, the overall inflation is not excessively high and does not significantly undermine the validity of our findings.

Nevertheless, having low power implies that the significant associations observed in our study may represent genes with an especially high impact on heat stress resistance. For example, *PRKG1* has already been confirmed as a heat stress-associated gene in other species, including humans. This strengthens the relevance of our findings. However, the functional validation of SNPs located in both genes is warranted to further elucidate the underlying physiological mechanisms.

Moreover, further studies with larger sample sizes are needed to verify our findings and eventually identify additional SNPs and candidate genes with lower effects on heat stress. Expanding the sample size in future research will enhance the robustness of our results and potentially uncover a broader spectrum of genetic factors involved in heat stress resistance.

## Conclusion

In this study, we identified significant associations between SNPs located in the *PDZRN4* and *PRKG1* genes and heat stress in cattle, providing important insights into the genetic basis of heat stress tolerance. Our findings contribute to the understanding of the physiological mechanisms underlying the heat stress response and provide potential genetic markers for selecting heat stress tolerant animals in breeding programs. Due to the limited sample size, further research is needed to validate our findings and identify genes with low to moderate effects.

## Statements & Declarations

### Funding

The publication is financed under the Leading Research Groups support project from the subsidy increased for the period 2020–2025 in the amount of 2% of the subsidy referred to Art. 387 (3) of the Law of 20 July 2018 on Higher Education and Science, obtained in 2019. This study was supported by China Agriculture Research System of MOF and MARA; The Program for Changjiang Scholar and Innovation Research Team in University (IRT 15R62); the National Agricultural Genetic Improvement Program (2130135).

### Competing Interests

The authors have no relevant financial or non-financial interests to disclose.

### Author Contributions

BC conceived and conducted the experiment. BC analyzed the results and wrote the manuscript in consultation with YW. All authors reviewed the manuscript, read and approved the final manuscript.

### Ethics approval

The data collection process was carried out in strict accordance with the protocol approved by the Animal Welfare Committee of the China Agricultural University. All experimental protocols were approved by the Animal Welfare Committee of the China Agricultural University. All methods are reported in accordance with ARRIVE guidelines (https://arriveguidelines.org) for the reporting of animal experiments.

## Notes

### Competing Interest Statement

The authors have declared no competing interest.

### Summary of Updates

QQ plot and genomic inflation factors have been calculated

